# Paracrine CSF1 signaling regulates macrophage migration dynamics towards ovarian cancer cells in a 3D microfluidic model that recapitulates *in vivo* infiltration patterns in patient-derived xenograft models

**DOI:** 10.1101/2022.09.27.509704

**Authors:** Alexis L Scott, Diana Kulawiec, Dorota Jazwinska, Ioannis K Zervantonakis

## Abstract

Ovarian cancer is the second most deadly gynecologic cancer in the United States, and tumorassociated macrophages in the ovarian cancer microenvironment are the most abundant immune cell type and are associated poor survival. Here, we utilize three-dimensional microfluidic assays to investigate the dynamics of macrophage infiltration towards ovarian cancer cells. Experimental results demonstrate that both ovarian cancer cell lines and patient-derived xenograft models promote the infiltration of macrophages into a 3D collagen type I extracellular matrix. Additionally, blocking CSF1 signaling reduced the number of recruited macrophages as well as migration speed, while macrophage recruitment was enhanced by addition of recombinant CSF1. We further demonstrated that results obtained with our microfluidic model are consistent with the recruitment of macrophages *in vivo* by patient-derived xenograft models, and that a xenograft model with high CSF1 expression showed an enhanced ability to recruit macrophages both *in vitro* and *in vivo*. These results highlight the role of CSF1 signaling in ovarian cancer, as well as the utility of microfluidic models in recapitulating the 3D ovarian cancer microenvironment.

## 1. Introduction

Ovarian cancer is the second most lethal gynecologic malignancy nationally, and the presence of alternatively activated macrophages in the tumor microenvironment has been associated with poor survival in ovarian cancer[1]. Macrophages in the tumor microenvironment may be recruited as circulating monocytes, which differentiate into tumor-associated macrophages (TAMs) upon entry. An array of paracrine signals derived from the tumor and stroma may influence the migration and phenotype of tumor-associated macrophages[2]. Namely, macrophage colony-stimulating factor (M-CSF/CSF1) has been shown to mediate the infiltration of macrophages in breast cancer[3], hepatocellular carcinoma[4], and melanoma[5]. CSF1 is also overexpressed in ovarian cancer and is associated with poor prognoses [6]. Therefore, inhibition of the CSF1 signaling axis presents a potential therapeutic approach to reduce tumor-promoting macrophages in the ovarian cancer microenvironment. Indeed, CSF1R inhibition (CSF1Ri) has been shown to decrease markers of M2-like tumor-promoting macrophages in 3D[7]. Furthermore, CSF1Ri delivery *in vivo* has been shown to deplete macrophages in glioblastoma as well as enhancing the infiltration of cytotoxic T cells and promoting survival[8]. However, the effect of CSF1 signaling on migration parameters such as the speed and infiltration distance of individual macrophages in a 3D ovarian cancer microenvironment is unknown. Therefore, novel approaches that can monitor the dynamics of cancer-macrophages in a precisely controlled microenvironment are necessary to dissect paracrine signaling mechanisms.

Other studies of macrophage recruitment in cancer have used culture systems that fail to accurately replicate the complexity of the tumor microenvironment. For example, cells cultured in 3D display differences in drug response, signaling, and morphology compared to 2D cultures[9,10]. To this end, we are employing a 3D physiologically relevant microfluidic platform to precisely control interactions between macrophages and tumor cells. This experimental design permits the establishment of chemokine gradients across the device, as well as live-cell imaging of macrophage chemotaxis. Previous studies have additionally used microfluidic devices to study the effects of interstitial flow and cytokine gradients on macrophage migration in the tumor microenvironment[11]. Using this platform, we have studied the dynamics of macrophage recruitment by ovarian cancer cells as mediated by CSF1. As CSF1 represents a key chemokine overexpressed in the ovarian tumor microenvironment[12], we tested the effects of small molecular inhibitors and blocking antibodies on macrophage infiltration towards ovarian cancer cells in a 3D matrix. Elucidating the mechanisms of macrophage recruitment in ovarian cancer may reveal druggable targets to block macrophage infiltration in ovarian cancer and improve patient outcomes.

## 2 Results

### 2.1 Microfluidic model of macrophage recruitment in ovarian cancer

The microfluidic device used to model macrophage recruitment was fabricated using molded polydimethylsiloxane (PDMS), which was adhered to glass coverslips via plasma bonding. The device consists of a central chamber containing ovarian cancer cells and collagen I, which is flanked by two fluidically independent side channels which are used for perfusing medium with suspended macrophages[13]. Ovarian cancer cells were engineered to express RFP (ID8H2BRFP, DF216-RFP, DF83-RFP) and were suspended at a density of 1e6 cells/mL in collagen. Following a 30-minute incubation to allow for the hydrogel to polymerize, macrophages were seeded in one of the two side channels at a density of 2e6 cells/mL (**Figure 1a**). To observe infiltration of macrophages into the collagen matrix, we utilized confocal microscopy, which demonstrated that macrophages are recruited into the device via migration through collagen (**Figure 1b, c**). Macrophages migrate in a 3D tumor microenvironment in response to cancer-derived chemoattractants, such as CSF1[12], IL-8 and CCL2,[11] and periostin[14]. As tumor secreted CSF1 is associated with poor prognoses in ovarian cancer[2], we specifically address the role of CSF1 in the recruitment of macrophages in the present work (**Figure 1d**).

**Figure 1:**
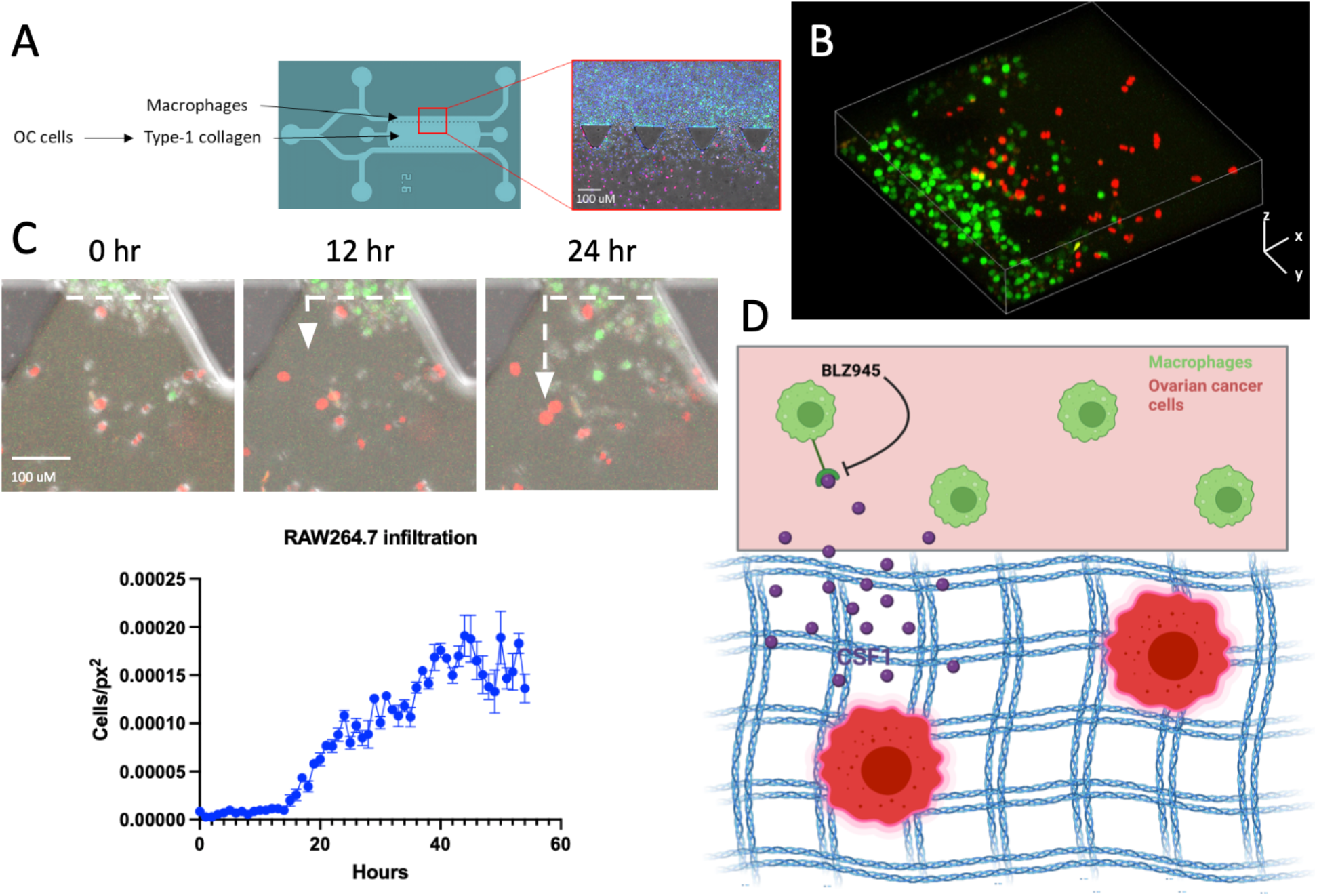
3D microfluidic modeling of ovarian cancer-macrophage interactions and macrophage infiltration. (A) Schematic of cell seeding in microfluidic device with ID8 ovarian cancer cells (red), RAW264.7 macrophages (green), and nuclei (DAPI). (B) 3D construction of confocal images demonstrating RAW264.7 (green) infiltrating into the ID8 (red)-containing gel after 2 days in microfluidic device. (C) Representative images of RAW264.7 infiltrating towards ID8 at 0, 12, and 24 hours after seeding. Dotted line represents gel-media interface. Quantified RAW264.7 cells per area within the gel region over time. Error bars represents SEM. (D) Schematic demonstrating the hypothesized effect of CSF1 and BLZ945 the modeled tumor microenvironment and subsequent macrophage recruitment.

### 2.2 Ovarian cancer secreted factors promote macrophage recruitment in 3D

To assess the ability of ovarian cancer cells to recruit macrophages in microfluidic devices, the number of RAW264.7 macrophages per area within microfluidic devices was analyzed with and without coculture with ID8 ovarian cancer cells. ID8 is a well-characterized model of ovarian cancer that was derived from C57/BL/6 ovarian epithelium. Specifically, ID8 is an aggressive model that establishes in vivo microenvironments that are infiltrated by macrophages[15]. Without tumor cells, the presence of macrophages within the gel region was 56 ± 6 × 10^−5^ cells/px^2^. The presence of tumor cells in the gel region increased macrophage infiltration to 93 ± 8 × 10^−5^ cells/px^2^ where statistical analysis (Student’s t test) revealed a significant difference (*p* = 0.02) (**Figure 2a, b**). Furthermore, we demonstrated that the number of recruited RAW264.7 positively correlates with the seeding density of ID8, and that treatment with ID8-conditioned media also enhances macrophage recruitment ∼2-fold (**Figure S1**). We additionally used a parallel 96-well plate-based collagen droplet assay to validate results obtained using microfluidic devices (Figure S2). Indeed, macrophages in monoculture infiltrated at 64 ± 3 cells/px^2^ in collagen droplets, while macrophages in coculture infiltrated at 19 ± 2 cells/px^2^ (*p* = 0.05) (**Figure 2a, c**). We additionally validated these results using mouse bone marrow derived macrophages (BMDMs) cocultured with ID8. BMDMs alone infiltrated at 3 × 10^−5^ ± 0.3 × 10^−6^ cells/px^2^, while coculture with ID8 enhanced BMDM infiltration to 19 ± 4 × 10^−5^ cells/px^2^ (*p* = 0.02) (**Figure 2d**).

**Figure 2:**
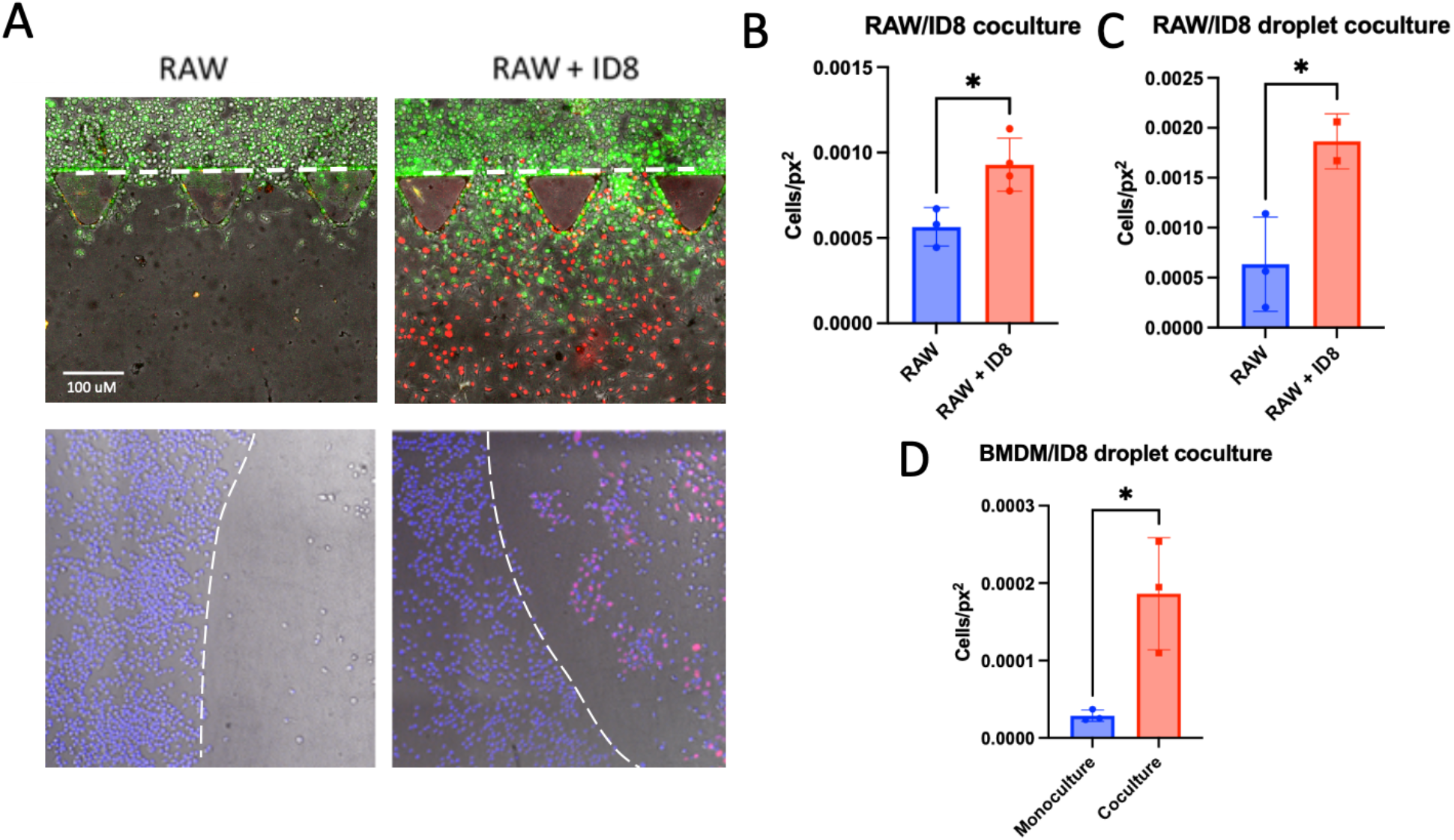
Ovarian cancer cells promote macrophage infiltration at endpoint. (A) Representative images 48 hours after seeding of microfluidic devices (top) or collagen droplets (bottom) containing either RAW264.7 macrophages in monoculture or with ID8 embedded in collagen. (B) Quantified RAW264.7 macrophages per area within devices in monoculture or cocultured with ID8 after 2 days. Unpaired t test (p = 0.0190). (C) Quantified macrophages per area in collagen droplets after 2 days. (D) Quantified BMDMs per area in collagen droplets after 2 days. Representative results from 3 devices per condition shown. Error bars represent SEM.

### 2.3 Ovarian cancer cell secreted factors enhance macrophage migration speed in 3D

Combining microfluidic systems with live-cell imaging allows for the simulation of dynamic factors in the tumor microenvironment such as migration speed[11,16]. As such, we utilized this approach to test the effect of tumor-secreted factors on macrophage migration speed in microfluidic devices (**Figure 3a**). When cultured alone, RAW264.7 macrophages migrated into the gel chamber at a speed of 68 ± 1.3 × 10^−5^ µm/sec. In coculture with ID8, the migration of RAW264.7 macrophages increased to 83 ± 4.50 × 10^−5^ µm/sec (*p* = 0.003, Student’s t test) (**Figure 3b, d**). We additionally used human PBMC-derived macrophages to validate these results when cocultured with human ovarian cancer cells. In this experiment, the human xenograft model DF216 was utilized. We found that PBMC-derived macrophages alone infiltrate at a speed of 14 ± 4 × 10^−5^ µm/sec, while coculture with DF216 enhanced recruitment speed to 15 ± 2 × 10^−5^ µm/sec (*p* = 0.05, Student’s t test) (**Figure 3c, e**).

**Figure 3:**
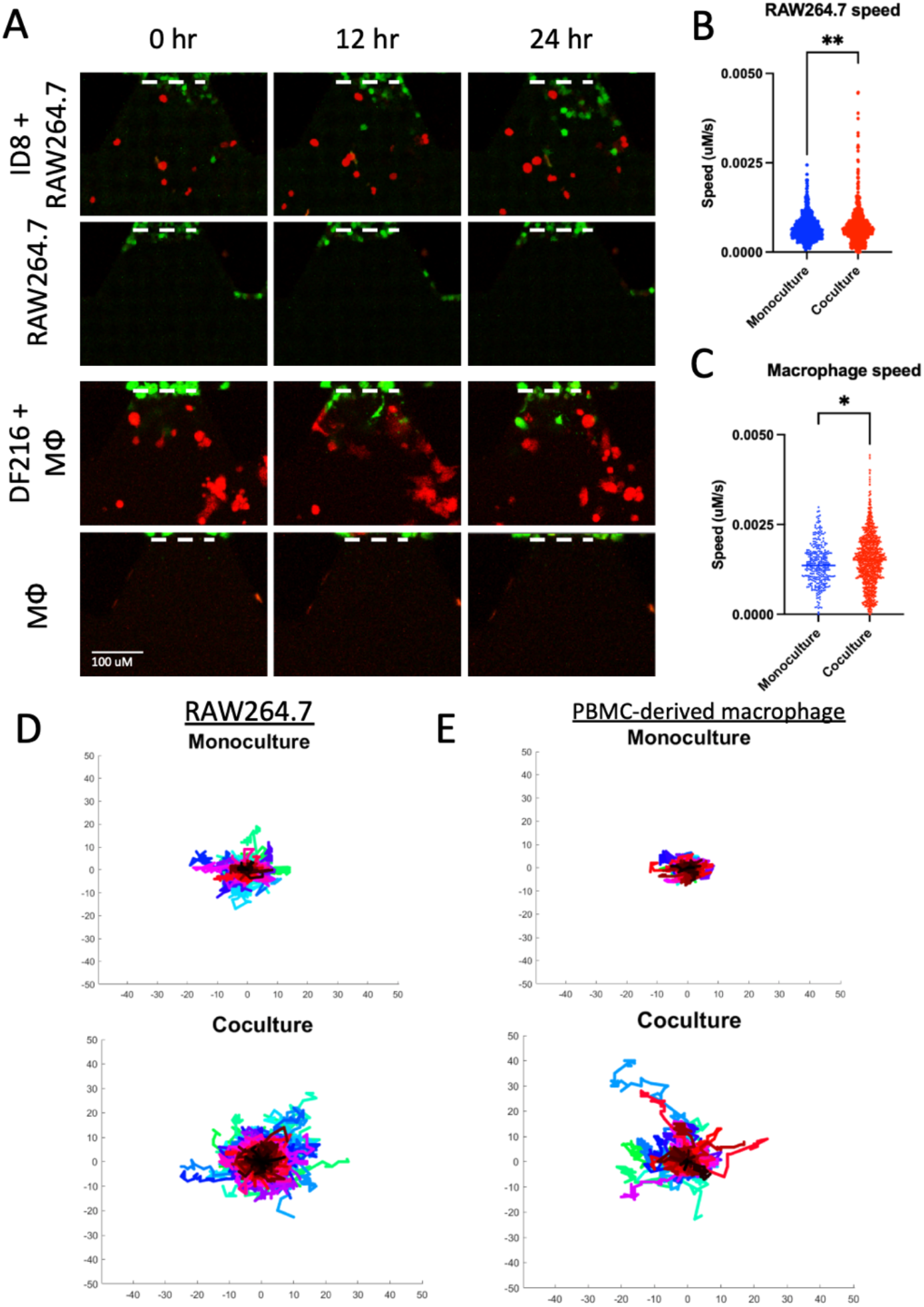
Time-lapse infiltration dynamics of recruited macrophages in microfluidic devices. (A) Representative images of RAW264.7 (green) and ID8 (red) or PBMC-derived macrophages (green) and DF216 (red) in microfluidic devices at 0hr, 12hr, and 2hr after macrophage seeding. (B) Speed of single RAW264.7 cell trajectories measured over 2 days. Representative results from 3 devices per condition displayed in which 6 interfaces per device were analyzed. (C) Speed of single PBMC-derived macrophages measured over 2 days. Representative results from 3 devices per condition displayed in which 6 interfaces per device were analyzed. Error bars represent SEM. (D) Wind-Rose plots displaying trajectories of RAW264.7 in monoculture or cocultured with ID8 in microfluidic devices over 2 days. (E) Wind-Rose plots displaying trajectories of PBMC-derived macrophages in monoculture or cocultured with DF216 in microfluidic devices over 2 days.

### 2.4 CSF1 signaling modulates macrophage recruitment speed

Next, we sought to examine the role of macrophage colony-stimulating factor (CSF1) signaling on the dynamics of macrophage recruitment in ovarian cancer using our microfluidic platform. CSF1 is essential for macrophage homeostasis and was identified for its ability to promote the expansion of macrophages from bone marrow progenitors[12,17]. As such, there is great interest in therapeutic strategies to hamper the activity of tumor-associated macrophages via blockade of CSF1 signaling. We therefore tested the effect of BLZ945, a potent CSF1R inhibitor, on macrophage migration speed. BLZ945 significantly reduced phospho-CSF1R staining in RAW264.7 macrophages, demonstrating the efficacy of the drug (**Figure 4a, b**). BLZ945 reduced macrophage infiltration speed from 19 ± 9 × 10^−5^ µm/sec to 14 ± 3 × 10^−5^ µm,/sec (*p* < 0.0001, Student’s t test) (**Figure 4c, Figure S3**). Additionally, BLZ945 reduced the path length of recruited macrophages from 23.8 ± 0.7 µm to 17.4 ± 0.5 µm (*p* < 0.0001, Student’s t test) (**Figure 4d, e**). BLZ945 also reduced the number of infiltrated macrophages in collagen droplets from 0.0024± 0.0003 cells/px^2^ to 0.00082 ± 0.0001 cells/px^2^ (*p* = 0.006, Student’s t test) (**Figure 4f**). To further validate these results, we also tested the effect of recombinant CSF1 as well as a CSF1 neutralizing antibody on macrophage recruitment. Addition of recombinant CSF1 to macrophages in monoculture increased the recruitment of macrophages by a factor of 3 ± 0.6 (*p* = 0.0481, Student’s t test). Once again, it was seen that coculture with ID8 significantly increased recruitment of RAW264.7 by a factor of 15.5 ± 2.6 cells/px^2^ (*p* = 0.0050, Student’s t test). Treatment of cocultures with an anti-CSF1 antibody reduced infiltration by 30% (*p* = 0.0147, Student’s t test) (**Figure S4**).

**Figure 4:**
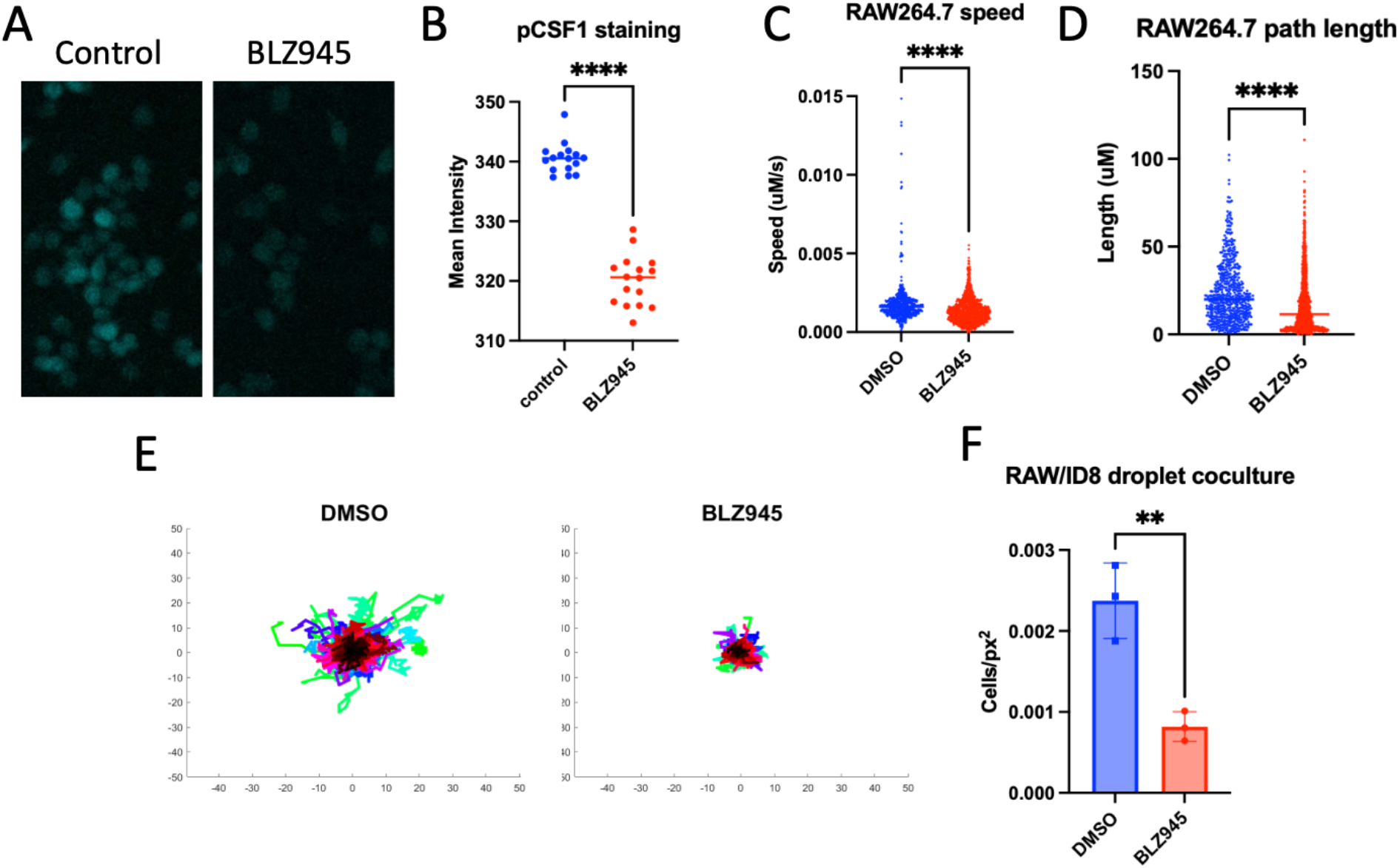
Effect of BLZ945 on macrophage infiltration dynamics. (A) Representative images of RAW264.7 macrophages treated with DMSO or BLZ945 stained with anti-pCSF1. (B) Quantification of mean fluorescence intensity from A. (C) Speed of single BLZ945 or DMSO-treated RAW264.7 cell trajectories measured over 2 days. (D) Path length of single BLZ945 or DMSO-treated RAW264.7 cell trajectories measured over 2 days. Representative results from 3 devices per condition displayed in which 6 interfaces per device were analyzed. (E) Wind-Rose plots displaying trajectories of DMSO or BLZ945-treated RAW264.7 over 2 days. (F) Number of RAW264.7 per area in collagen droplets with treatment of BLZ945 or DMSO. (F) Normalized infiltration of RAW264.7 macrophages with treatment of recombinant CSF1 and normalized infiltration of RAW264.7 in coculture with ID8 with treatment of anti-CSF1 neutralizing antibody. Error bars represent SEM.

These results demonstrate the critical role of the CSF1 signaling axis in modulating macrophage migration properties in a 3D matrix.

### 2.5 Patient-derived cells with high *CSF1* expression recruit more macrophages *in vivo* and in microfluidic devices

To further explore the role of CSF1 signaling in the 3D ovarian cancer microenvironment, we sought to use patient-derived ovarian cancer cell lines established from xenograft models. From a panel of xenograft models, DF216 was selected as a model for high *CSF1* expression and DF83 was selected as a model for low expression. Using bulk RNA-seq from DF216 and DF83 xenografts, it was observed that *CSF1* expression in these models was 315 and 10 counts per million respectively (**Figure S5**). As DF216 has a significantly higher *CSF1* expression compared to DF83, we next characterized macrophage recruitment by these cells *in vivo*. Indeed, a higher fraction of F4/80 positive cells was observed in DF216 tumors compared to DF83 tumors, reflecting the trend of differential *CSF1* expression (**Figure 5a, b**). Next, the macrophage recruiting capabilities of DF216 and DF83 spheroids were tested using microfluidic devices (**Figure 5c**). Reflecting macrophage recruitment *in vivo*, DF216 recruited significantly more macrophages into the gel region of devices compared to DF83 (**Figure 5d**). To account for differential growth rates between DF83 and DF216, we additionally measured the number of RAW264.7 cells in the gel regions of microfluidic devices as a factor of %RFP, once again revealing that DF216 more potently recruit macrophages compared to DF83 (**Figure S5**). As macrophage recruitment in microfluidic devices reflects results obtained using the same tumor model *in vivo*, these results demonstrate the physiological relevance of using microfluidic models to study macrophage infiltration.

**Figure 5:**
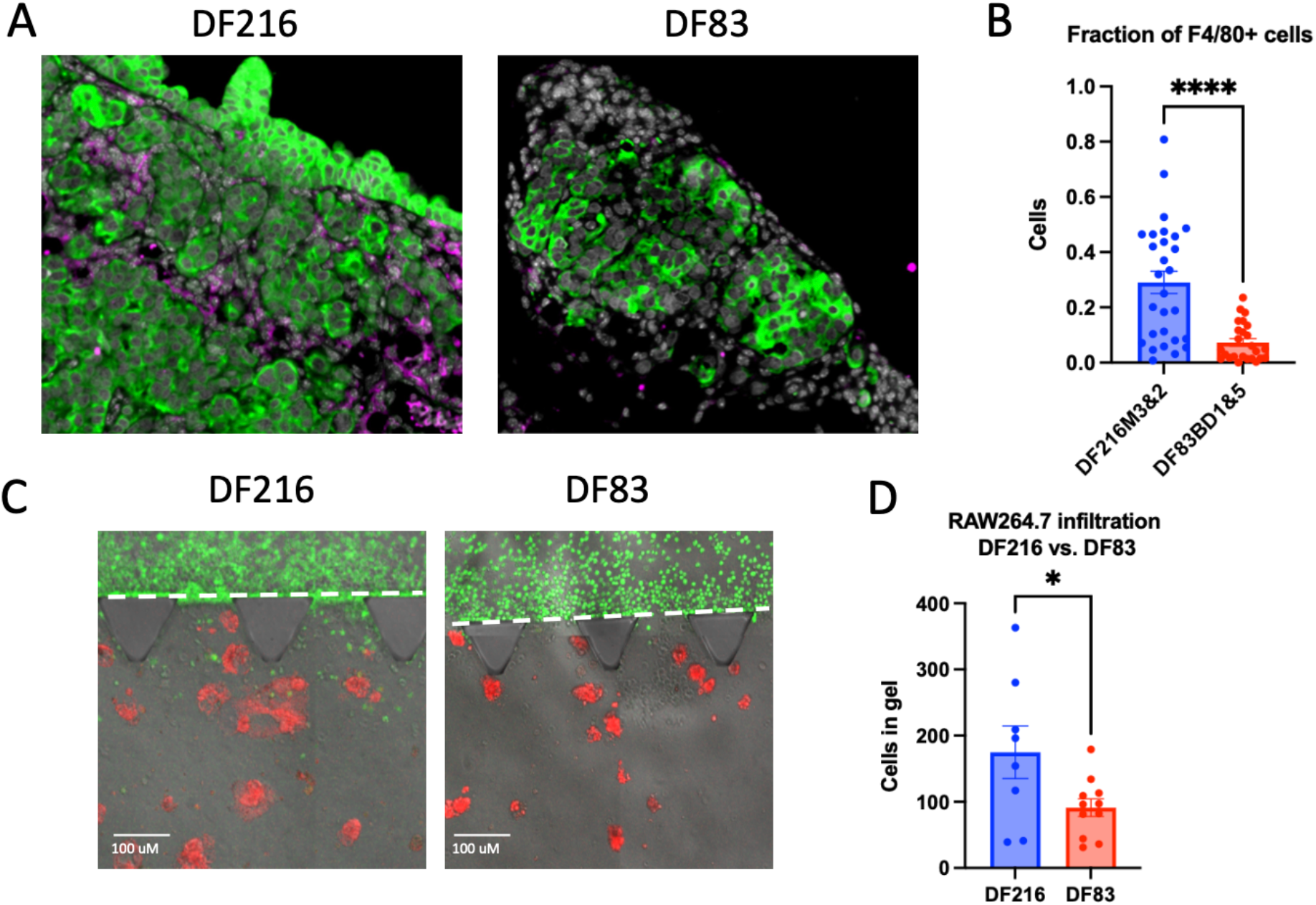
3D culture systems align with in vivo results. (A) Pan-cytokeratin (green) and F4/80 (magenta) staining of DF83 and DF216 xenograft tumor models. (B) Quantified fraction of F4/80 cells per field of DF83 and DF216 xenografts. (C) Representative images of RAW264.7 macrophages infiltrating towards DF83 or DF216 in microfluidic devices. (D) Number of RAW264.7 cells in gel region of microfluidic devices with DF216 or DF83 after 2 days. Error bars represent SEM.

## 3. Discussion

In summary, by combining 3D cell culture in microfluidic devices with live-cell confocal imaging, we demonstrated the recruitment of macrophages towards ovarian cancer cells as modulated by CSF1. We further highlighted the physiological relevance of this platform for studying the macrophage recruitment and the CSF1 signaling axis by the agreement between our microfluidic system and *in vivo* data using patient-derived tumor models. These results indicate that microfluidic culture systems can readily recapitulate the tumor microenvironment of ovarian cancer as modulated by CSF1, reinforcing the utility of these systems in studying clinically relevant models.

In ovarian cancer, as well as breast cancer[18–20], Hodgkin’s lymphoma[21], and non-small cell lung cancer[22], macrophage infiltrates are associated with poor clinical outcomes. Given the tumor-promoting functions displayed by macrophages in ovarian cancer, there lies significant potential in utilizing CSF1-targeted therapies to treat the disease. For example, macrophages mediate resistance to chemotherapy and radiotherapy via STAT3 activation in tumor cells[23,24]. Macrophage migration additionally supports tumor progression and metastasis, as matrix remodeling by macrophages allows tumor cells to migrate via “tracks’’ created by macrophages in the extracellular matrix[25]. Tumor-associated macrophages also frequently possess tumor-promoting immunomodulatory functions, such as suppressing T-cell activation via the consumption of metabolites, inducing regulatory T cells via cytokine and chemokine secretion, and generally inducing immune suppression via expression of checkpoint ligands[23,26]. Therapeutic targeting of CSF1 signaling may therefore present a strategy to skew the tumor microenvironment of ovarian cancer away from an immunosuppressive state. However, macrophages may also take part in immunosurveillance against cancer cells, as they are capable of phagocytosis of malignant cells and antigen presentation, which stimulate the proliferation of T-cells[27]. As such, the application of methods to study the dynamic interactions between ovarian cancer cells and macrophages is critical.

Using a microfluidic system, we first demonstrated that ovarian cancer cells enhance the number of recruited macrophages. We additionally validated these results using a parallel 3D collagen droplet assay. These results are similar to migration indices observed in transwell coculture assays with ovarian cancer cells and macrophages[14]. Furthermore, previous *in vivo* studies have shown that ID8 tumor models can be characterized by enrichment of macrophages in tumor nodules[15]. The recruitment of macrophages by ovarian cancer cells in our model is therefore reflective of previous *in vivo* and *in vitro* studies. Previous microfluidic studies have also demonstrated an enhancement of macrophage recruitment speed by other tumor-secreted factors (e.g. CCL2 and IL8 [11]). We additionally utilized patient-derived ovarian cancer cells to compare results obtained with our microfluidic system to *in vivo* macrophage responses. We demonstrated that primary cancer cells with a higher expression of *CSF1* more potently recruit macrophages both *in vivo* and in a microfluidic model. Furthermore, the results obtained using our microfluidic system are in agreement with previous studies, as Lu *et al*. demonstrated that breast cancer cell lines with greater CSF1 secretion recruit more macrophage *in vivo* and promote greater macrophage migration in a scratch assay[30].

As CSF1 signaling is implicated in the survival and chemotaxis of macrophages, we additionally sought to use our microfluidic model to assess the contribution of tumor secreted CSF1 to macrophage recruitment in ovarian cancer. In the ovarian cancer microenvironment, tumor cells are a major secretor of CSF1 [28]. As such, targeting CSF1 signaling is a promising strategy for treating ovarian cancer by ablating infiltration of tumor-promoting macrophages. Indeed, CSF1R inhibitors have shown clinical promise in ovarian cancer, as CSF1R blockade has been shown to reduce the number of M2 macrophages in ovarian tumors as well as decrease ascites volume[29]. In this study, we tested the effect of BLZ945, a potent inhibitor of CSF1R, on the dynamics of macrophage recruitment in a microfluidic model of ovarian cancer. We observed that BLZ945 reduced the speed and path length of recruited macrophages in a microfluidic model as well as the total number of recruited macrophages in a collagen droplet assay. These results align with previous work demonstrating that BLZ945 reduces abundance of tumor-associated macrophages in a murine model of ovarian cancer[24]. While previous studies utilizing 3D cell culture have established that CSF1R inhibition via BLZ945 reduces M2-like macrophages in 3D models[7], this work is the first to demonstrate the effect of BLZ945 directly on macrophage migration through a collagen matrix.

Altogether, we have demonstrated the utility of a microfluidic device to recapitulate macrophage recruitment in ovarian cancer. We specifically showed that the CSF1 signaling axis plays a vital role in the recruitment dynamics of infiltrating macrophages in ovarian cancer. While our microfluidic model accurately represents the spatial interactions between tumor cells and macrophages in ovarian cancer, incorporation of additional cell types in the model such as cancer-associated fibroblasts could further recapitulate the ovarian cancer microenvironment. The adaptability of this system easily allows for the incorporation of additional cell types and biologic variables. Further developments of 3-dimensional assays to recapitulate paracrine tumor-stroma interactions will allow for discovering novel therapeutic targets in a physiologically relevant context.

## 4 Experimental Section

### Cell Culture

ID8 (provided by MIT Swanson Biotechnology Center) were cultured in DMEM supplemented with 10% fetal bovine serum (FBS) and 1% penicillin/streptomycin. High grade serous ovarian cancer patient-derived cells DF83 and DF216 (described in Liu et al 2016) and were also cultured in DMEM supplemented with 10% fetal bovine serum (FBS) and 1% penicillin/streptomycin. RAW264.7 were purchased from (ATCC) and cultured in DMEM supplemented with 10% fetal bovine serum (FBS) and 1% penicillin/streptomycin. Mouse monocytes were isolated from long bones of female FVB/Blc6 mice according to Amend, Valkenburg, & Pienta and cultured in 20 ng/mL recombinant mouse M-CSF[31]. CD14+ monocytes were isolated from normal human PBMCs (ATCC PCS-800-011™) using CD14+ microbeads and LS columns (Miltenyi, USA) Monocytes were cultured in Blood Cell Culture Medium (iXCells, USA) supplemented with 100 ng/mL recombinant human M-CSF.

### Microfluidic Infiltration Assay

Collagen was prepared using 20 µL PBS, 10 µL NaOH, 60 µL H20, and 110 µL Type-1 rat tail collagen (Corning, USA). Ovarian cancer cells were suspended 1e6/mL and seeded in the central chamber of the microfluidic device. Collagen was polymerized for 30 minutes at 37C. After polymerization, DMEM (Corning, USA) w/ 10% fetal bovine serum and 1% penicillin/streptomycin was added to the inlet ports and aspirated through the channels of the device. Following a 30-minute equilibration period, macrophages were seeded at a concentration of 2e6 cells/mL in DMEM (Corning, USA) w/ 10% fetal bovine serum and 1% penicillin/streptomycin. Imaging was performed with a Zeiss LSM700 Confocal microscope and images were analyzed using Nikon Elements.

### BLZ945 treatment of microfluidic cocultures

RAW264.7 or PBMC-derived macrophages were suspended in media supplemented with 5 µM BLZ945 (MedChemExpress, USA) and seeded at 2e6 cells/mL in microfluidic devices as described previously. Media with BLZ945 was also added to the opposite channel of the device.

### Collagen Droplet Infiltration Assay

Collagen was prepared using 20 µL PBS, 10 µL NaOH, 60 µL H20, and 110 µL Type-1 rat tail collagen (Corning, USA). Ovarian cancer cells were suspended 1e6 cells/mL in 3 µL collagen droplets. Collagen was polymerized for 30 minutes at 37C. Macrophages were added at a concentration of 25,000 cells/well in DMEM with 10% fetal bovine serum and 1% penicillin/streptomycin. To study the effect of CSF1 signaling in droplet cultures, wells were treated with either 5 µM BLZ945 or 20 ng/mL recombinant mouse M-CSF at the time of macrophage seeding. After 48 hours, the plate was fixed with 4% paraformaldehyde in PBS and stained overnight with Hoechst. Imaging was completed with a Zeiss LSM700 Confocal microscope.

### Image Analysis

Microfluidic and collagen droplet cultures were imaged at day 2 and analyzed with NIS-Elements software. Raw images with GFP or cellTracker labeled macrophages were binarized using the bright spots function, and an ROI containing the collagen-containing section of each image was drawn manually, with the functional output being the number of labeled objects per drawn ROI area.

### Cell Motility

Images were analyzed using NIS-Elements software with Tracking module. For time-lapse images, microfluidic devices were imaged every 15 minutes for 48 hours. 3 gel-media interfaces per device were selected for time-lapse analysis based on interface integrity. Raw FITC images were binarized using the bright spots tool, and the tracking module was used to assign each migrating macrophage to a track and compute dynamic features. Cell migration speed was calculated as:

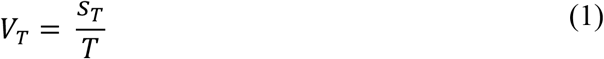

where *S*_*T*_ = length and *T* = total time lapsed. Length was calculated as:

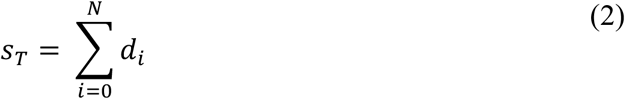

where *N* = number of segments and *d*_*i*_ = segment length.

### Statistics

Results are displayed as the mean ± standard error. The Student’s t-test was used to evaluate comparisons between groups in Figures 2b-d, 3b-c, 4b-c, 4e-f, 5c, and 5e. Statistical significance was considered to be *p* < 0.05. Statistical tests were performed with Prism (Version 9.0, GraphPad Software).

---

**Supplementary Figure 1:**
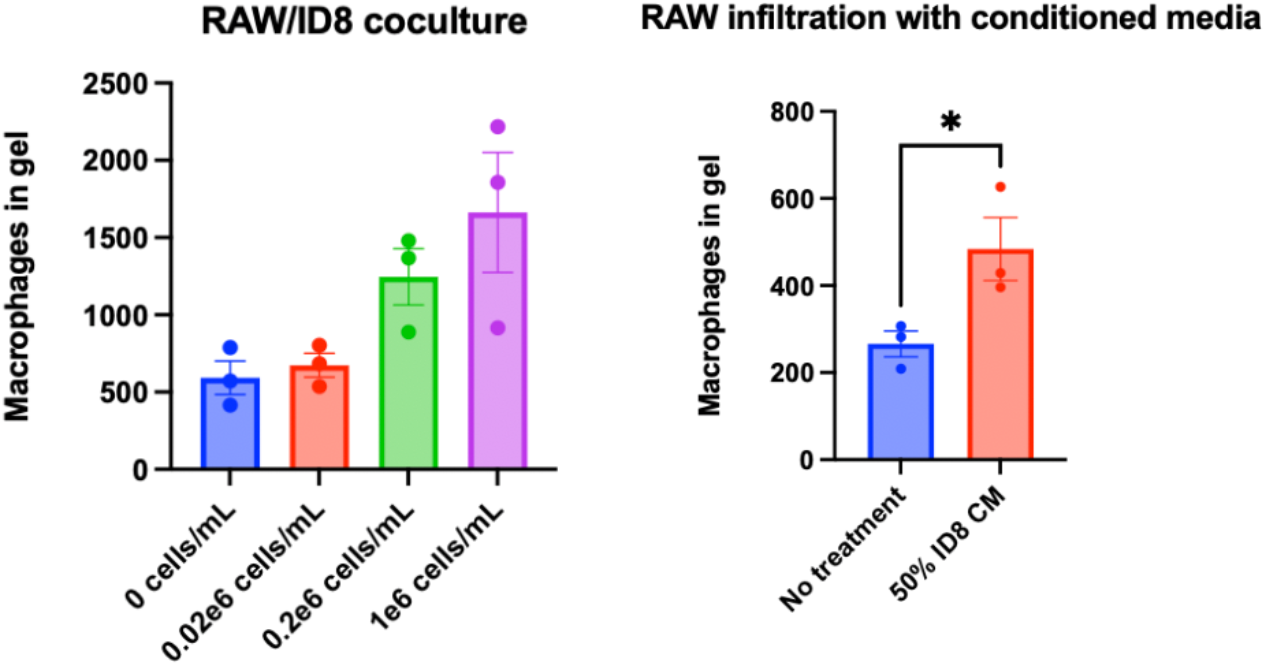
Cancer cell density and cancer conditioned medium. (A) Increasing cancer cell density resulted in highr numbers of recruited RAW264.7 macrophages in the 3D collagen gel. (B) RAW264.7 macrophage infiltration following treatment with 50% control and 50% ID8 conditioned medium in the 3D collagen gel assay.

**Supplementary Figure 2:**
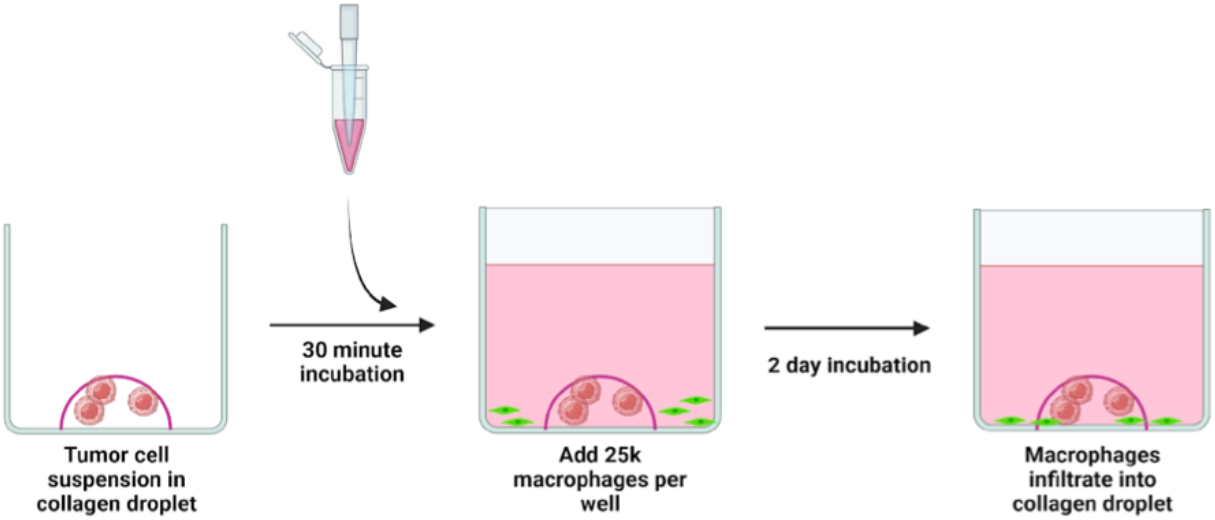
Schematic of 3D collagen gel assay. Macrophage infiltration is monitored 48hr following seeding on top of a 3D collagen type I gel with embedded cancer cells.

**Supplementary Figure 3:**
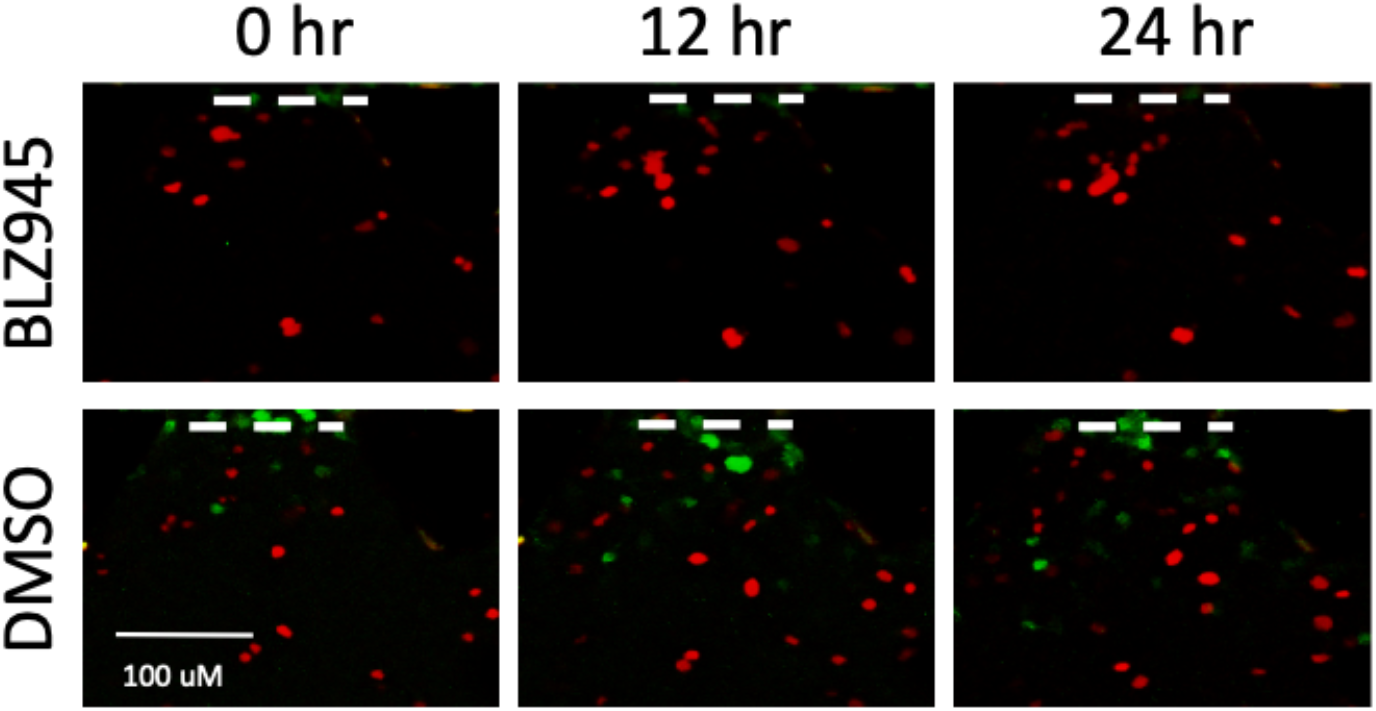
BLZ945 treatment in microfluidic devices. Time-lapse images of devices (ID8 and RAW264.7 coculture) treated with BLZ945

**Supplementary Figure 4:**
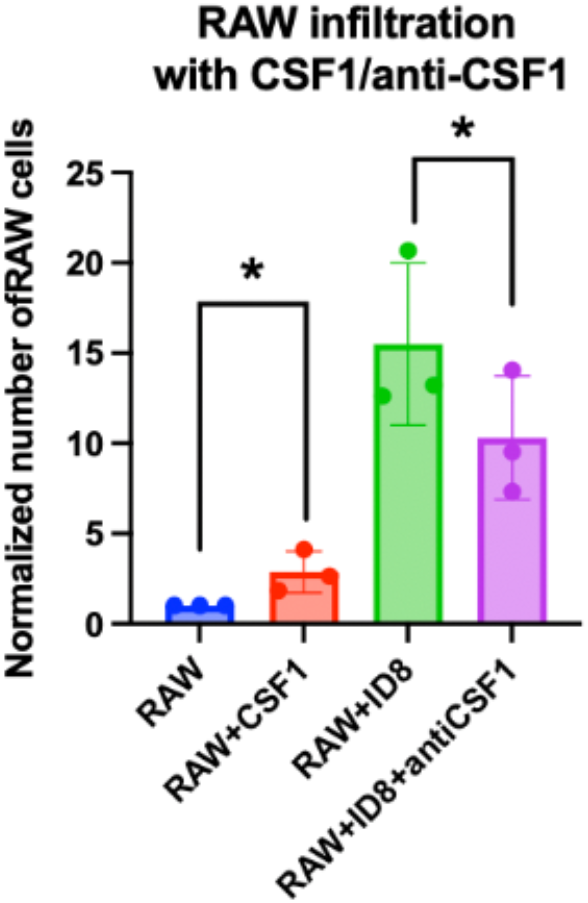
Exogenous CSF1 and anti-CSF1 blocking treatment in 3D collagen gel assay. RAW264.7 infiltration was assessed following treatment with CSF1 (20ng/ml) or coculture with ID8 +/-anti-CSF1 blocking.

**Supplementary Figure 5:**
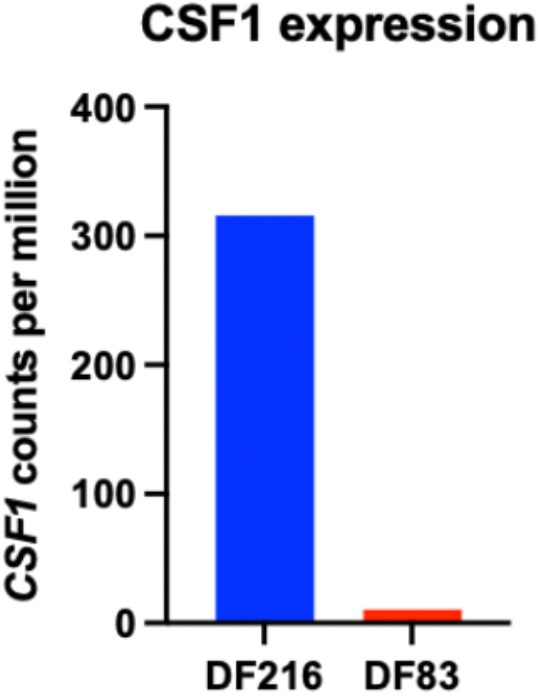
Expression of CSF1 transcript levels in two patient derived xenograft models.

